# Sensitive Chromosomal Translocation Quantitation from Amplicon Sequencing Using Primer-Anchored Statistical Translocation Analysis (PASTA)

**DOI:** 10.64898/2026.07.23.740402

**Authors:** Ellen Schmaljohn, Olalekan Usman, Christian Brommel, Kyle J. Kinney, Nathan White, Giandomenico Turchiano, Thomas Osborne, Andrea Sánchez-Peña, John Sterrett, Rolf Turk, Garrett Rettig, Ashley M. Jacobi, Morgan Sturgeon, Gavin L. Kurgan

**Author notes:** These authors contributed equally to this work and share first authorship (Ellen Schmaljohn, Olalekan Usman).

## Abstract

Chromosomal translocations are rare structural rearrangement outcomes of genome editing, requiring analytical frameworks that combine high quantitative accuracy with performant sensitivity and specificity. Amplicon sequencing offers a scalable means to detect rare rearrangements with ultra-deep targeted sequencing, but existing methods often rely on heuristic thresholds or ad hoc normalization steps that limit reproducibility and have unknown analytical performance. Here, we present a computational tool we call PASTA (Primer-Anchored Statistical Translocation Analysis), using a count-based differential-event statistical framework to quantify and statistically confirm translocation junctions from targeted amplicon sequencing data. Comparison of this method to ddPCR demonstrates that quantitation is highly accurate, and outperforms other NGS-based methods even when randomized adapter chemistry is not present in amplicon sequencing structures. To measure analytical performance, we create a benchmarking dataset for measuring chromosomal translocation analysis performance with frequencies ranging from 1% to sub-0.01%, and demonstrate that the method can detect frequencies down to 0.01% with >75% sensitivity when sufficient read depth is present. Taken together, this work demonstrates using amplicon sequencing with PASTA as a bioinformatics analysis tool is a solution for translocation detection in amplicon sequencing genotoxicity assessments, enabling identification of rare genome rearrangements in both research and preclinical applications

## Introduction

Genome editing technologies such as CRISPR-Cas systems induce targeted double-strand breaks (DSBs) or single-strand breaks (SSBs) to achieve precise genetic modification. These engineered DNA lesions are repaired by endogenous pathways, most prominently non-homologous end joining (NHEJ) and other alternative end-joining mechanisms, that can produce both accurate and error-prone repair outcomes^1^. Although the intended repair product in therapeutic gene editing work often restores reading frames or introduces designed sequence changes, unintended events such as large deletions, chromosomal rearrangements, and complex genome alterations have been repeatedly observed^2–5^. Because these aberrant outcomes may carry genotoxic risk, quantitatively characterizing the full spectrum of repair products, including rare chromosomal translocations, is essential for pre-clinical safety assessment and mitigation of adverse outcomes in therapeutic genome editing.

Chromosomal translocations represent a particularly concerning class of genomic rearrangements in the context of genome editing because of their potentially unpredictable and difficult-to-characterize therapeutic consequences. By juxtaposing genomic elements that are not normally linked, translocations can alter gene expression, disrupt coding sequences or regulatory elements, generate fusion transcripts, and, in some cases, contribute to malignant transformation or other pathogenic phenotypes^6–9^. While most translocation events generated during genome editing are thought to be deleterious and quickly face progressive dilution of affected cells over time (especially with *ex vivo* expansion), some have been shown to persist^10,11^. In the context of genome editing, both DSB and SSB have been shown to induce chromosomal translocations, although their frequencies vary substantially across target sites, modalities, and experimental systems^12–14^. In general, DSB-generating editors have been shown to produce translocations at higher frequencies than SSB-based editors, likely due to the increased availability of free DNA ends capable of aberrant rejoining^13^. However, translocation formation remains highly dependent on guide RNA specificity, off-target activity, chromosomal architecture, and DNA repair pathway utilization^10,12,13^. Given their potential safety implications, translocation analyses have recently been recommended for routine genome-editing characterization in therapeutic packages generating DSBs^15^. Consequently, determining reliable methods for quantifying the true prevalence and long-term persistence of editing-induced translocations has become an important challenge in the assessment of genome-editing safety.

Several approaches have been developed to quantify gene editing outcomes, with next generation sequencing (NGS) emerging as the gold standard due to its ability to sensitively and accurately capture the diversity of sequence alterations created at on- and off-target sites. Analytical frameworks have been developed and demonstrated high performance in profiling insertions, deletions and other DSB-associated repair signatures with high reproducibility across target sites using NGS data^16,17^. However, for chromosomal translocation quantitation, the gold-standard has traditionally relied on low-throughput techniques such as karyotyping and targeted droplet digital PCR (ddPCR)^18,19^. While both technologies have demonstrated generally high specificity and quantitative precision, they have sensitivity limitations at sub-0.1% events and are challenging to multiplex. Recent work has improved some of these limitations through multiplexing of ddPCR^20^, Optical Genome Mapping^21^, or single-cell workflows like Directional Genomic Hybridization (dGH)^22^, though important limitations to sensitivity, throughput, and quantitative resolution still exist. Furthermore, many of these methods require equipment and/or workflows that are not routine for organizations developing gene editing therapeutics. Thus, as genome editing advances toward therapeutic applications, higher-throughput, statistically robust and sensitive methods that fit conventional workflows are ideal for routinely assessing the genotoxic potential of editing agents.

NGS-based sequencing methods have become widely available and have largely expanded capabilities for translocation detection through advances in both library preparation strategies and computational algorithms. These different methods can be classified into a few main groups: uni-directional anchoring and amplification strategies, hybridization capture, and traditional amplicon sequencing coupled with specialized detection algorithms. For example, UdiTas^23^ and CAST-seq^14^ represent uni-directional anchoring strategies that enable partially-biased discovery of translocations and rearrangements by leveraging specialized library construction to capture aberrant junctions at locations originating from known target locations (e.g. the on-target). However, detection scope is currently restricted to events where the on-target is a participating partner and its workflow is thus not currently scalable for large-panel safety testing. Alternative analytical approaches such as CRISPECTOR apply statistical modeling to amplicon sequencing data to infer translocation events, but the method lacks built-in methods to fully exploit the statistical strength provided by experimental replicates and mechanisms to convert counts to translocation frequencies^24^. Collectively, all these methods still lack comprehensive benchmarking for identifying and quantifying well-described translocations across different sequence space as well, meaning important metrics like analytical sensitivity, specificity, and quantitative precision are mostly uncharacterized. Thus, there remains a need for benchmarked, scalable, and statistically rigorous NGS-based approaches that can reliably quantify low-frequency translocations across diverse experimental contexts.

Advances in RNA-seq differential expression analysis have produced a diverse ecosystem of statistical frameworks that share several core principles essential for accurate interpretation of count-based genomic data^25,26^. These methods, including widely used algorithms built upon generalized linear models, empirical Bayes shrinkage, and robust normalization strategies have undergone extensive benchmarking across large RNA-seq datasets to evaluate their performance under varying levels of sequencing depth, dispersion, and experimental complexity^27–29^. The analytical strengths of these frameworks have led to their extension beyond classical gene expression profiling into applications requiring highly sensitive detection of rare genomic events, such as low frequency variants in ultra-deep sequencing workflows, including somatic variant classification^30^. Their ability to model overdispersed count data, incorporate replicate structure, and provide principled statistical inference makes them naturally suited for translocation detection from amplicon sequencing data. This is especially valuable because true junction counts must be distinguished from stochastic amplification artifacts and index hopping events^31–33^. As a result, RNA-seq differential expression algorithms may provide a mature, rigorously tested statistical foundation for developing methods to quantify rare genome-editing outcomes.

To meet regulatory expectations for pre-clinical safety testing, analytical frameworks should also be accompanied by comprehensive benchmarking that defines their performance boundaries and operational reliability. Parameters such as input genomic copy number, sequencing depth, replicate structure, library conversion rates, and assay design can substantially influence event detection rates, as has been shown in previous works^34^. Amplicon architecture can have a particularly important effect on translocation detection in amplicons sequencing based on whether P5/P7 adapter sequences are randomized or fixed. Fixed adapters to forward and reverse primers can bias capture of primer-pair combinations and lead to systematic loss of specific translocation events, potentially reducing confidence in quantitative results^10,24^. As has been demonstrated in fields like somatic variant calling, benchmarking datasets and controlled experimental designs are necessary to characterize detection limits, sensitivity, specificity, and reproducibility. Applying similar rigor to gene editing translocation analysis (where translocations can occur at sub-1% frequencies regularly) will ensure that NGS-based methods are fit-for-purpose in genotoxicity risk assessment and can support accurate, reproducible, and regulator-ready-evaluation of genome editing safety.

To address these shortcomings and maximize use of existing amplicon sequencing data for pre-clinical safety assessment, we created a computational method we term Primer-Anchored Statistical Translocation Assessment (PASTA). By treating each putative forward–reverse primer pair as an independent “event feature” within a negative binomial modeling framework implemented in DESeq2, the method robustly accounts for sampling variance, dispersion, and technical noise inherent in multiplexed PCR-based assays. By iteratively testing different mathematical quantitation methods against well-established orthogonal methods like ddPCR, we determine an ideal method to calculate translocation frequency from amplicon sequencing and demonstrate improved performance compared to other methods, such as CAST-seq. By developing an ultra-deep sequencing translocation benchmarking dataset, we demonstrate that this approach enables highly quantitative estimation of translocation frequencies, achieving sensitive detection down to translocation frequencies of 0.01% while maintaining stringent control of false positives. Finally, we apply this method to draw general biological insights on features leading to translocation occurrence, providing general guiding principles for further supporting early de-risking and identification of events.

## Results

### Primer-Anchored Statistical Translocation Analysis (PASTA) Algorithm

Following gene editing, translocations or other large re-arrangements can be amplified from sequencing data and with primers identified, used to create count matrices for statistical testing of significantly enriched events above typical background sequencing events, such as index hopping as has been previously shown^24^ (Figure 1A). However, the ability to detect translocations in amplicon sequencing can be hindered dependent on the library preparation methods utilized. Fixed adapter chemistry, such as rhAmpSeq, have sequencing adapters only associated with the forward or reverse primer in a pre-specified manner, which may lead to cases where certain primer pairs (such as two forward primers) cannot participate in bridge amplification for detection by sequencing (Figure 1B). Randomized adapter chemistry, such as TQ-rhAmpSeq^35^, has sequencing adapters paired with any forward or reverse primer, which may enable more complete detection by ensuring that all pairs of forward and reverse primers can form sequenced products (Figure 1C).

**Figure 1.**
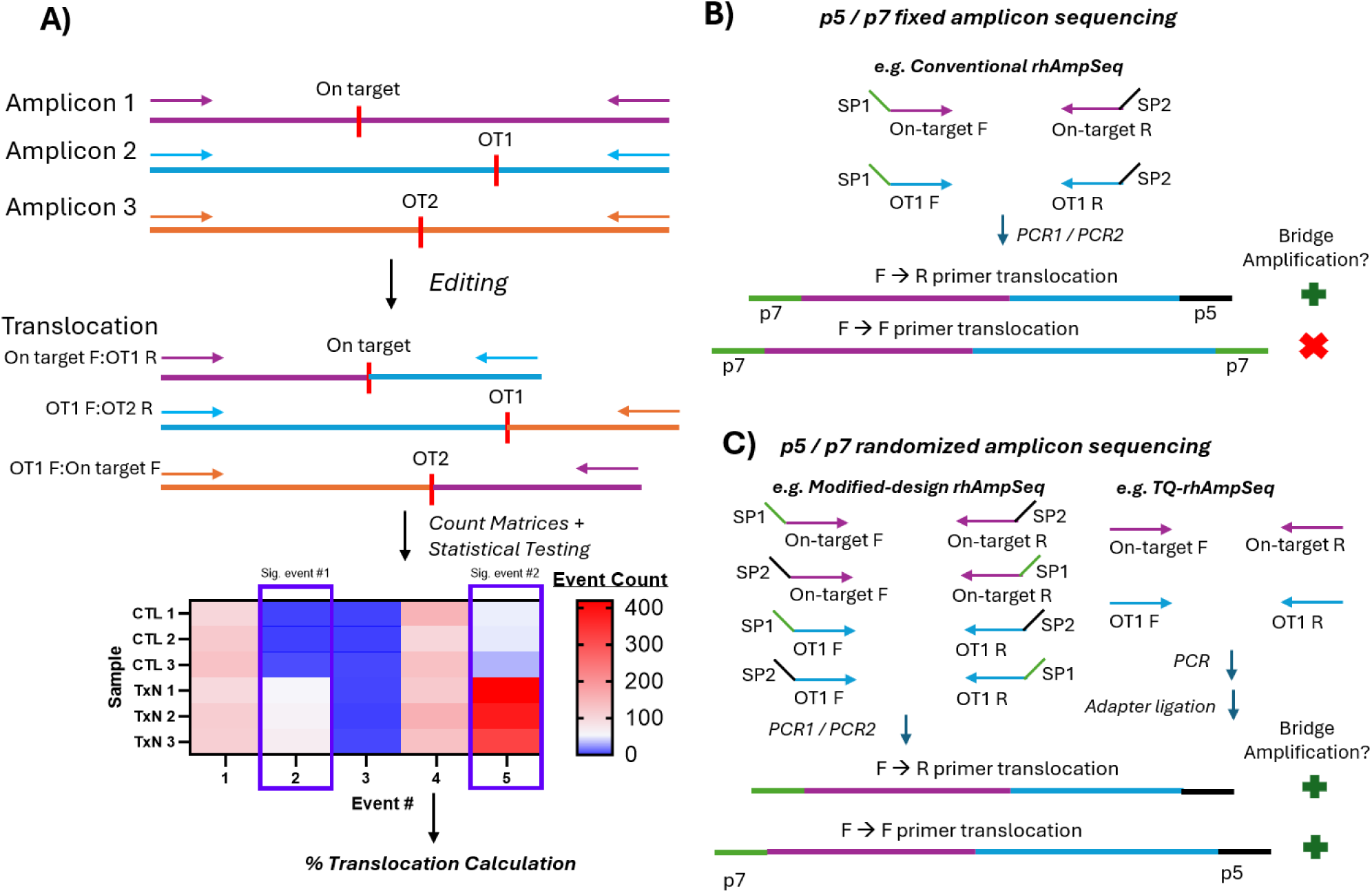
Primer Anchored Statistical Translocation Analysis (PASTA) algorithm diagram. **A)** An example shows how multiplexed amplicon sequencing following a gene editing cut site (red line) can be used to detect events of translocated amplicon using counts of primer pairs identified in sequencing. Amplicon structures can be organized to capture different types of translocation events with examples shown for what can be captured with **B)** fixed and **C)** randomized adapter amplicon structures.

To create the PASTA algorithm, we rationalized that the tool should: 1) function off of raw sequencing data as a starting point to enable re-analysis of other datasets (independent of other CRISPR or gene editing analysis algorithms), 2) have methods to handle highly variable experimental structures for statistical analysis indicative of real world limitations (n = 1 vs n > 1), 3) function in a standardized software environment (e.g. Docker) without assumptions of specific workflow management tools (e.g. Nextflow) to prevent any software variability caused by implementation or infrastructure incompatibility, and 4) handle variable amplicon sequencing structures (e.g. fixed P5/P7 adapters versus random P5/P7 adapters).

The resulting algorithm workflow is shown (Supplementary Figure 1). Initially, primers are either provided explicitly or inferred based on amplicon sequencing structures, and reads are aligned to appropriate amplicon ends with mismatch tolerance. Following alignment, count frequencies are calculated from individual primer pairings that were observed in all treatment and control samples, within respective R1/R2 FASTQ files. To enable count normalization and performance testing at variable read depth, we built per-assay downsampling capabilities to enable optional read depth normalization. To handle variable replicate structures, two different statistical analysis methods were enabled, a hypergeometric test with multiple-testing correction for N=1 experimental structures (assuming a single paired treatment and control sample; hereafter referred to as ‘hypergeometric’), and a count-based negative binomial modeling framework for all other replicate structures (assuming at least two or more paired treatment and control samples; hereafter referred to as ‘DESeq2’ since the DESeq2^25^ package was used as the basis for performing this). After significance testing, quantitative translocation frequencies are inferred based on the frequencies of the event observed and the frequencies of the involved primers in other intended and unintended pairings. Following this, heuristic filters are applied to flag or filter events that could either be indicative of outliers that can cause false positives (e.g. insufficient read depth for intended frequencies) or have significant deviations that warrant additional review or removal (see Methods). The workflow then outputs the translocation events being observed as well as their estimated quantitative frequencies and marks the completion of the PASTA workflow.

### Characterization of Quantitative Accuracy Relative to ddPCR

Accurate absolute quantification of the frequencies of translocations is necessary to understand the cellular burden of these genomic arrangements in a translational context. To determine an optimal equation for translocation frequency quantification from significant event counts in NGS data, ddPCR assay data from previous work^35^ was re-analyzed as a truth set for the translocation frequencies of measuring a single translocation (off-target_1_:on-target) and a cumulative set of translocations between an on-target and its off-targets (on-target:off-target_all_) under four variable editing conditions resulting in translocation differences of a promiscuous CCR5 gRNA across 20 different loci (n = 4 single translocation datapoints; n = 4 cumulative translocation datapoints) (Figure 1A). Importantly, the same genomic DNA used for ddPCR was also in parallel prepared in this previous work for sequencing using TQ-rhAmpSeq and CAST-seq^35^. Translocation frequencies quantified via ddPCR ranged from 10.6% to 0.3% translocation across editing conditions (Figure 2). The accuracy of different PASTA translocation quantitation equations varied, with some equations trending to overestimate ddPCR translocation values (Equation 1 and 2) and others trending to underestimate ddPCR translocation values (Equation 3 and 4). The worst equation (Equation 1) overestimated translocation frequency by an average log2 fold-change of 1.5 ± 0.7 from ddPCR derived values (Supplementary Figure 2). Using the average count of the forward and reverse primers involved in the translocation event as the denominator yielded the best agreement with ddPCR without trending to underestimate translocation frequency, with an average log2 fold-change of 0.0 ± 0.6. Thus, the “Equation 2” calculation was used for the PASTA algorithm to transform count data to interpretable translocation frequency values (Supplementary Figure 2).

**Figure 2.**
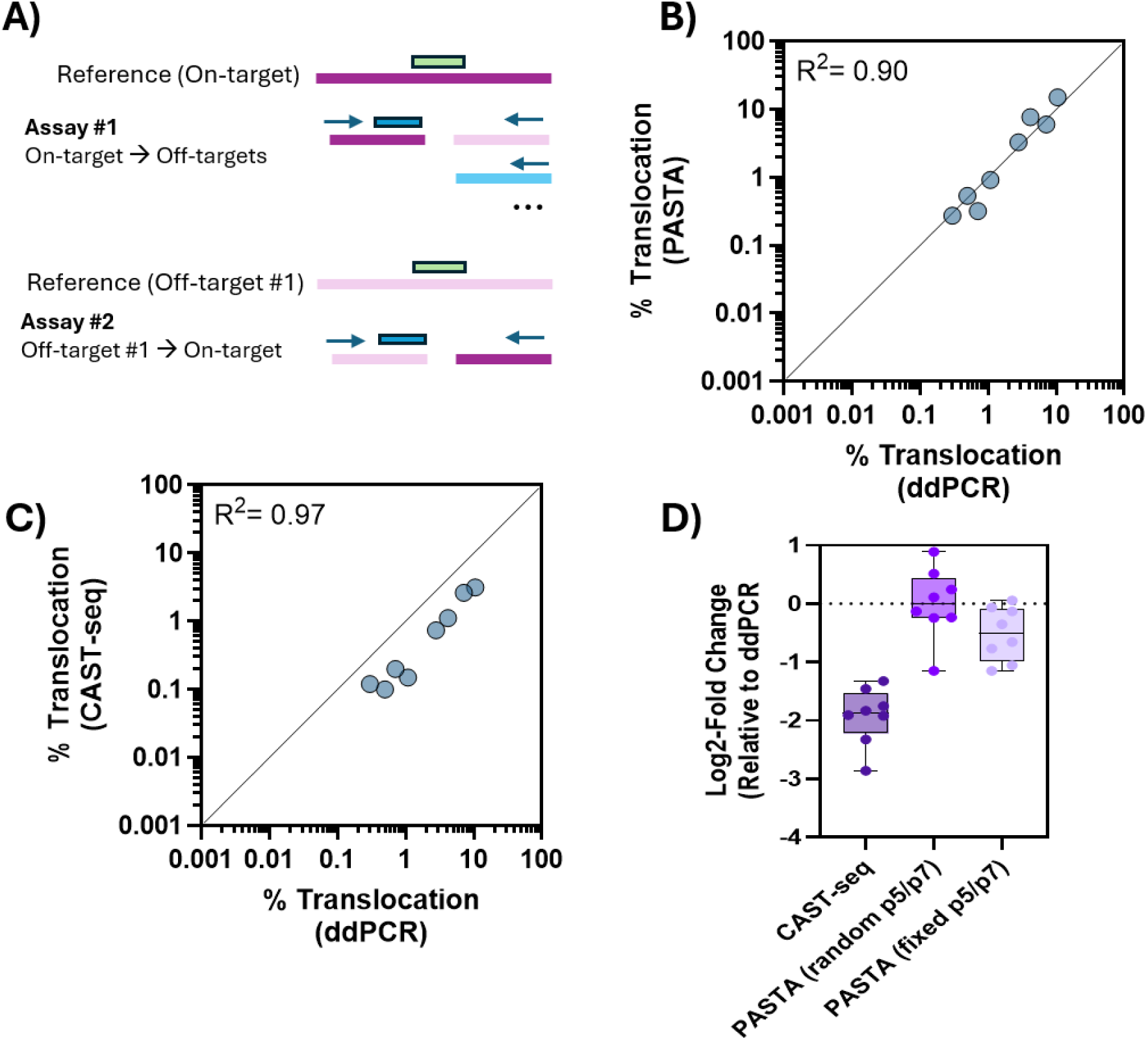
Quantitative precision of PASTA and CAST-seq using ddPCR as ground-truth. **A)** ddPCR assay design concept showing probe position for the reference (green) vs targeted translocation event (blue) for an assay designed to capture a cumulative frequency of the on-target to all off-targets (#1) and for an assay capturing a single off-target translocated to the on-target. Quantitative frequencies of translocations from NGS were compared to ddPCR derived values (two assays; four treatments; n=8 per method) for both **B)** PASTA and **C)** CAST-seq. **D)** For all NGS methods the calculated fold change from ddPCR was calculated and shown as a box plot. Fixed p5/p7 PASTA was derived from random p5/p7 PASTA data by removing translocations that would be plausible in the fixed adapter chemistry.

To compare the quantitative performance of randomized adapter amplicon sequencing (TQ-rhAmpSeq) using PASTA to other methods, we compared to orthogonally generated CAST-seq data prepared from the same gDNA in parallel. PASTA using randomized adapter amplicon sequencing yielded highly correlative data with ddPCR (R^2^= 0.90), and median log2 fold-change from ddPCR derived frequencies of 0.0 ± 0.6 (Figure 2B; Figure 2D). CAST-seq generated translocation frequencies were also highly correlative with ddPCR (R^2^=0.97), however CAST-seq consistently underestimated translocation frequency with a median log2 fold-change from ddPCR-derived frequencies of −1.9 ± 0.5 (Figure 1C; Figure 1D). To test the quantitative accuracy of amplicon sequencing and PASTA with fixed adapter chemistry, we next investigated the performance of PASTA while removing all translocations that would not have been possible under fixed adapter chemistry (forward primer-forward primer pairs and reverse primer-reverse primer pairs). Using fixed adapter chemistry, the quantitative accuracy of amplicon sequencing with PASTA dropped from random adapter chemistry performance due to missed events, but still outperformed CAST-seq in total quantitative accuracy for the events with a log2 fold-change of −0.5 ± 0.5 (Figure 1D).

To further ensure that the algorithm was able to correctly distinguish events that should be present only in randomized adapter chemistry (e.g. primers paired in F/F or R/R conformations) vs fixed adapter chemistry (e.g. primers paired in F/R conformations), we further re-analyzed paired rhAmpSeq and TQ-rhAmpSeq data from previously published work using the same CCR5 gRNA under conditions leading to variable off-target frequencies^35^. In this case, data generated with fixed adapter chemistry (rhAmpSeq) should generally not produce any significant events in orientations it cannot support (F/F and R/R), whereas TQ-rhAmpSeq should produce significant events. Quantified read depth was similar between the fixed and randomized adapter chemistry datasets with a range of 2 to 4 million reads per sample (Supplementary Figure 3A). Most amplicons had similar read depth between the datasets, though some assays were better sequenced in one dataset than the other, with a Spearman correlation of 0.44 (Supplementary Figure 3B).

Significant F/R translocations that should be identified in both amplicon sequencing chemistries had similar event numbers quantified in both datasets (Supplementary Figure 3C), while significant F/F and R/R translocation pairs were only identified in randomized adapter chemistry datasets, as expected (Supplementary Figure 3D). These results collectively show that following quantitation method optimization, both fixed and random adapter amplicon sequencing can be readily used to infer both individual and cumulative translocation frequencies with improved accuracy compared to an alternative library preparation and analysis method like CAST-seq.

### Characterization of Analytical Specificity Performance

To characterize the analytical specificity of amplicon sequencing with PASTA, we performed a readily available fixed adapter amplicon sequencing chemistry (rhAmpSeq) using a fixed genomic DNA input (100ng; ∼30,000 haploid genomes) to generate a dataset for testing that should have no translocations. To create an appropriate dataset that should have no translocations but could also be used for indel detection validation (Schmaljohn et al., in preparation), 23 distinct genomic loci were edited in single gRNA transfections in biological triplicate. The resulting gDNA was pooled to create a fixed concentration of each event (1% indels), and the pooled samples were serially diluted 3 orders of magnitude across a concentration range from 1% to 0.01% (Supplementary Figure 4A). Since the edited gDNA pooled are 23 distinct on-targets and not related by sequence homology, this should represent a multiplexed dataset with no possible translocations, and thus any significant events are false positives (Supplementary Figure 4A). Statistical tests also often gain confidence in deriving significant events from count data as count frequencies increase, leading us to hypothesize that method specificity may be impacted at variable read depths. To determine the impact of read depth on performance, we performed ultra-deep sequencing and downsampled read depth per assay to test deviations in specificity at variable read depth. This analysis was performed with both the hypergeometric method (n = 1 paired treatment/control per comparison) and DESeq2 method (n = 3 paired treatment/control per comparison) to quantify specificity differences between the statistical testing methods (Supplementary Figure 4A).

Characterization of the gene edited dataset showed, as intended, each on-target gRNA had high frequencies of on-target editing ranging from 28% to 98% indels in K562 cells, and that the unedited control samples had indel activity approaching the frequency of noise at 0.09% indels (Supplementary Figure 4B). Sequencing of the specificity dataset generated a coverage per assay ranging from 142,000x to 2,735,000x with a median of 708,000x, enabling us to check the effects of downsampled coverage on method performance from 1,000x to 500,000x (Supplementary Figure 4C). Following translocation classification, both the hypergeometric and DESeq2 methods showed 100% specificity down to read depths of 10,000x coverage per assay (Figure 3A). As read depth increased from 50,000x to 500,000x, the specificity of the hypergeometric testing method decreased from 85.8% to 71.5% (Figure 3A). Specificity for the DESeq2 method however remained high, with specificity remaining at 100% even at 500,000x coverage (Figure 3A). This analysis demonstrates that using the DESeq2 statistical testing method in PASTA while leveraging biological replicates increased specificity and is the preferred method of analysis for reducing false positives from translocation analysis.

**Figure 3.**
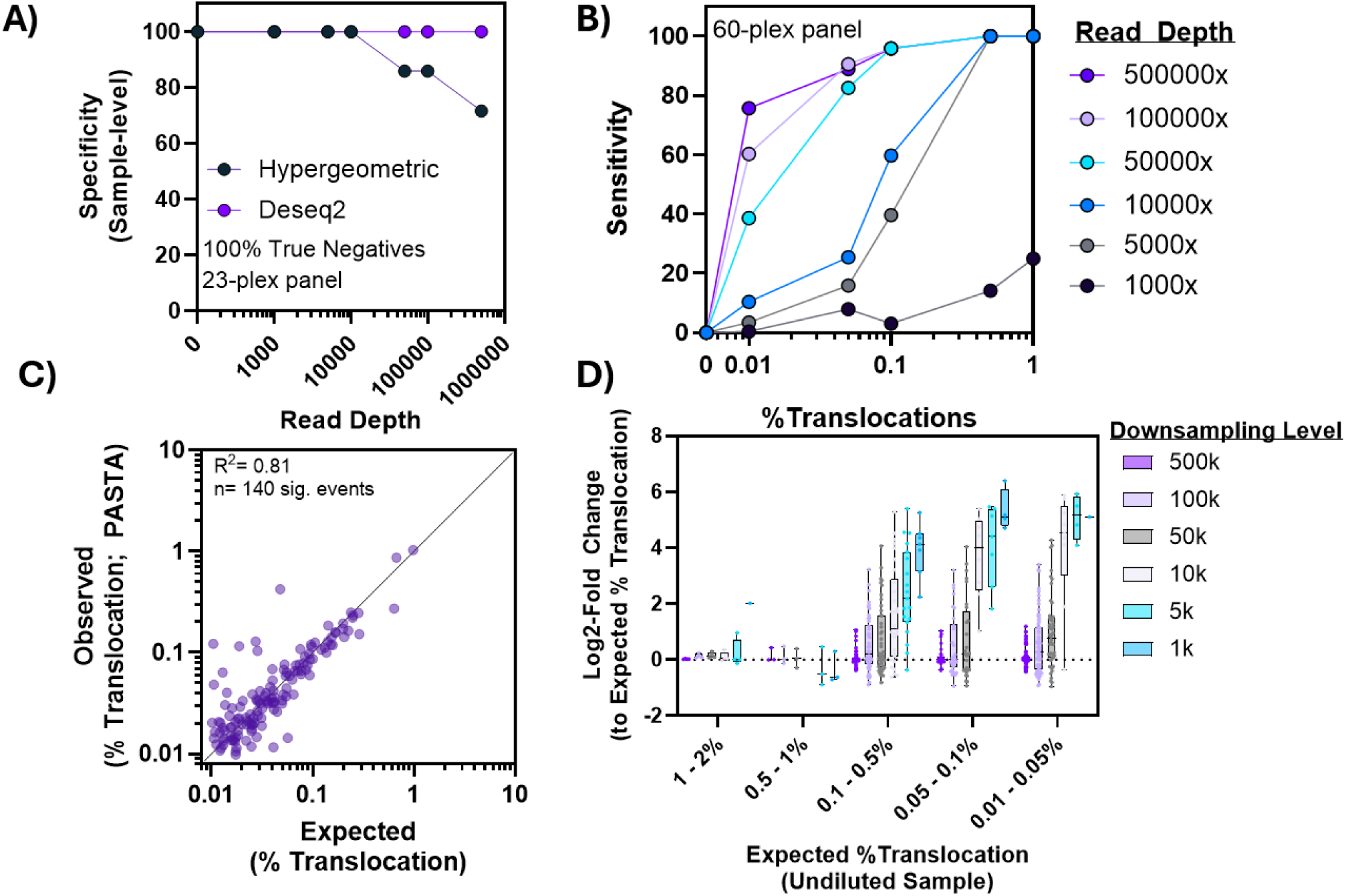
Sensitivity and specificity of PASTA statistical methods at variable read depth. **A)** Edited K562 gDNA samples (n = 22) were sequenced with a twenty-three multiplexed primer set of genomic targets that were unrelated to one another to test the specificity of the workflows. In these samples, any significant events thus were a true positive, and negatively impacted specificity. **B)** Edited HEK293-Cas9 samples using a LAG3 site 9 gRNA were serially diluted in biological triplicate (n = 3 treatment / control per dilution) after having true positive translocations identified in the undiluted condition. Sensitivity was measured as a fraction of the total available true positive events that should be present based on each dilution per downsampling level (500,000x to 1,000x). **C)** Measured translocation frequency of all significant translocations with expected and observed frequencies > 0.01% (n = 140; range from 1% to 0.01%) compared to what would be expected in the 50% dilution sample. **D)** Quantitative precision dependence on appropriate read depth was measured as log2 fold-change from the expected frequency in the undiluted sample originally characterized.

### Characterization of Analytical Sensitivity Performance

To characterize the analytical sensitivity of amplicon sequencing with PASTA, we again performed a readily available fixed adapter amplicon sequencing chemistry (rhAmpSeq) using a fixed genomic DNA input (100ng gDNA; ∼30,000 haploid genomes) (Supplementary Figure 5A). Since, to our knowledge, a ground-truth dataset of a highly diverse set of translocations at sub-10% translocation frequencies is not readily available, we opted to create our own by characterizing a gRNA’s off-targets and performing a serial dilution. To create a dataset suitable for sensitivity analysis, a well-characterized promiscuous guide RNA from previous work^12^, *LAG3* site 9, was used to edit stably expressing SpCas9 HEK293 cells in biological triplicate. Cells were edited and translocations characterized in a non-diluted state across a previously developed 60-plex multiplex amplicon sequencing panel targeting the on-target and 59 off-targets spread across 4 orders of magnitude of off-target editing^12^. The indel editing frequencies of these different selected sites varied greatly from 75% indels at the on-target down to 0.01% indels at the lowest significant off-target, with a total of 35 significantly edited sites (Supplementary Figure 5B). Furthermore, the indel frequencies observed were highly related to the Severity Bin of each off-target, with indel frequencies and significant off-target number decreasing as Severity Bin increased (Supplementary Figure 5C). Quantification of read depth showed that a median of 467,000x to 515,000x coverage was obtained across assays (Supplementary Figure 5D).

To determine the list of sites to be monitored for sensitivity analysis, the DESeq2 PASTA method was run on the panel of on- and off-target sites and all significant sites were labeled as true-positives, with the respective starting frequency recorded as ground-truth. This was rationalized to be appropriate since it was previously demonstrated that using the DESeq2 method with biological triplicate paired treatment:control samples resulted in 100% specificity, and the frequencies of PASTA for individual translocations closely mirrored ddPCR frequencies (Figure 2; Figure 3A). This resulted in detection of 179 unique translocations spanning from 0.01% to 2.0% initial frequencies to be true positive sites for sensitivity analysis (Supplementary Figure 6A). Following this, we serially diluted the genomic DNA at the following edited:unedited concentrations 50%, 10%, 5%, 1%, and 0.5% and characterized sensitivity for detecting these true positive translocations at variable read depths and absolute frequencies. The translocation frequencies occurring were binned as generalized frequencies to measure sensitivity in as follows: 0.01% < x ≤ 0.05%, 0.05% < x ≤ 0.1%, 0.1% < x ≤ 0.5%, 0.5% < x ≤ 1.0%, 1.0% < x ≤ 2.0%. Characterization of the total number of events (n) in each translocation events per bin ranged from 5 to 259 true positive events following dilution (Supplementary Figure 6B).

Since statistical tests lose the ability to significantly identify events as count frequencies decrease, we similarly sought to determine the impact of read depth on sensitivity using the recommended DESeq2 statistical analysis method. To this end, we downsampled read depth per assay to test deviations in sensitivity at variable read depth throughout the diluted samples (Supplementary Figure 5E). Furthermore, we hypothesized that since the number of hypotheses being tested rapidly expands with increasing amplicon panel sizes, that this could interfere with the ability to derive significance at lower amplicon read counts. Upon modeling how the hypothesis number could be impacted *in silico*, we quantified up to 3,600 or 21,240 hypothetical pairings for fixed and randomized adapter chemistry in a 60-plex amplicon panel, respectively (Supplementary Figure 7A). In cases where very large hypothesis numbers exist, filtering out non-relevant hypotheses is typically necessary to maintain high sensitivity following multiple-testing correction in biological data^25,36^.

Upon performing the initial sensitivity analysis, it was observed that sensitivity quickly dropped to 0% across all dilutions after read depth fell below 50,000x coverage, suggesting that multiple testing correction issues could be occurring at the higher multiplexing level (Supplementary Figure 7B). Measuring the count frequency of non-significant events in the treatment population demonstrated that non-significant events often had a median count frequency between 1-3 events, regardless of dilution (Supplementary Figure 7C). To test if reduction of non-relevant hypotheses could improve performance, we tested implementation of a count filter prior to statistical testing to remove events with less than 3 events in the treated population. Measurement of the number of hypotheses being tested prior to this filter at the 10,000x coverage dilution was 815 hypotheses, as compared to 219 hypotheses after this filter (Supplementary Figure 7D). This was shown to decrease the number of hypotheses tested by approximately 4-fold from the original unfiltered dataset, which we propose as the default to ensure more robust behavior using the PASTA tool while minimally impacting any real events (Supplementary Figure 7D).

Following this optimization, we re-characterized the sensitivity across the dilution series and different read coverage downsampling levels. This demonstrated that sensitivity as high as 75.6% was possible for detecting 0.01% translocation frequencies with 500,000x coverage (Figure 3B). As may be expected, sensitivity decreased as read depth decreased, though >95% sensitivity was still possible for detecting events down to 0.1% with 50,000x coverage (Figure 3B). Importantly, implementation of the minimum event filter increased performance at lower read depth subsampling levels from 0% sensitivity all the way to 100% sensitivity for detecting events >0.5% translocation frequencies with 10,000x coverage, demonstrating the impact of this optimization (Supplementary Figure 7; Figure 3B). By quantifying the observed frequencies of translocation detection compared to expected in the 50% dilution sample, a highly linear relationship was observed (R^2^= 0.81) between expected and observed frequencies all the way down to 0.01% (Figure 3C). Characterization of the impact of read depth in achieving high quantitative precision in the undiluted samples also demonstrated highly differential requirements for characterizing translocation frequencies to a median of 2-fold from the true frequency, ranging from 1,000x to 50,000x to quantitate a translocation to this resolution (Figure 3D). Taken together, this demonstrates that PASTA can sensitively and reproducibly detect true positive translocation events. Furthermore, we find the method has a highly linear range of detection to 0.01% and elucidates guidance on read depth requirements for quantitating different frequencies of translocations.

### Characterization of Relationships Between Translocation and Indel Editing

To better understand how editing information relates to translocation propensity and frequencies, we evaluated the translocations generated in the LAG3 site 9 gRNA with additional meta-data for this gRNA derived from previous nomination data^12^ and our own confirmation results. Determining empirical relationships between these meta-data would enable better triaging of early risk at nomination stages, and better expectations for supporting significant findings of targets following confirmation.

We first evaluated nomination propensity relationships to translocation propensity. In accordance with the UNCOVERseq nomination method, we binned the off-targets that we interrogated according to off-target nomination propensity using Severity Bins and quantified the number and frequency of translocations derived from each group. As might be expected, the highest frequency of translocations were derived from translocation events involving the on-target site, with an average translocation frequency of 0.25% derived from 21 separate events (Figure 4A). Furthermore, as the nomination frequency decreased, the average translocation frequency decreased from 0.14% with an average of 16.1 translocations per off-target in Severity Bin 1 down to 0.0% in Severity Bin 5 where no translocations were observed (Figure 4A). Between both translocation partners, generally high nomination frequencies were observed as well, with an average nomination frequency between partners of 81% relative to the on-target nomination frequency. When looking at the differences between each translocation partner nomination frequency, translocation partners generally had at least one high nomination frequency off-target, with a minimum nomination frequency within a pair of 22% the on-target (Figure 4B). Furthermore, on average, the more lowly nominated off-target in the pair was generally high as well with an average frequency of 55% the on-target within the lower nominated target within the translocation pair (Figure 4B). This information largely suggests that nomination frequencies from UNCOVERseq can be used to determine high risk translocation sites, especially when both sites have high frequencies.

**Figure 4.**
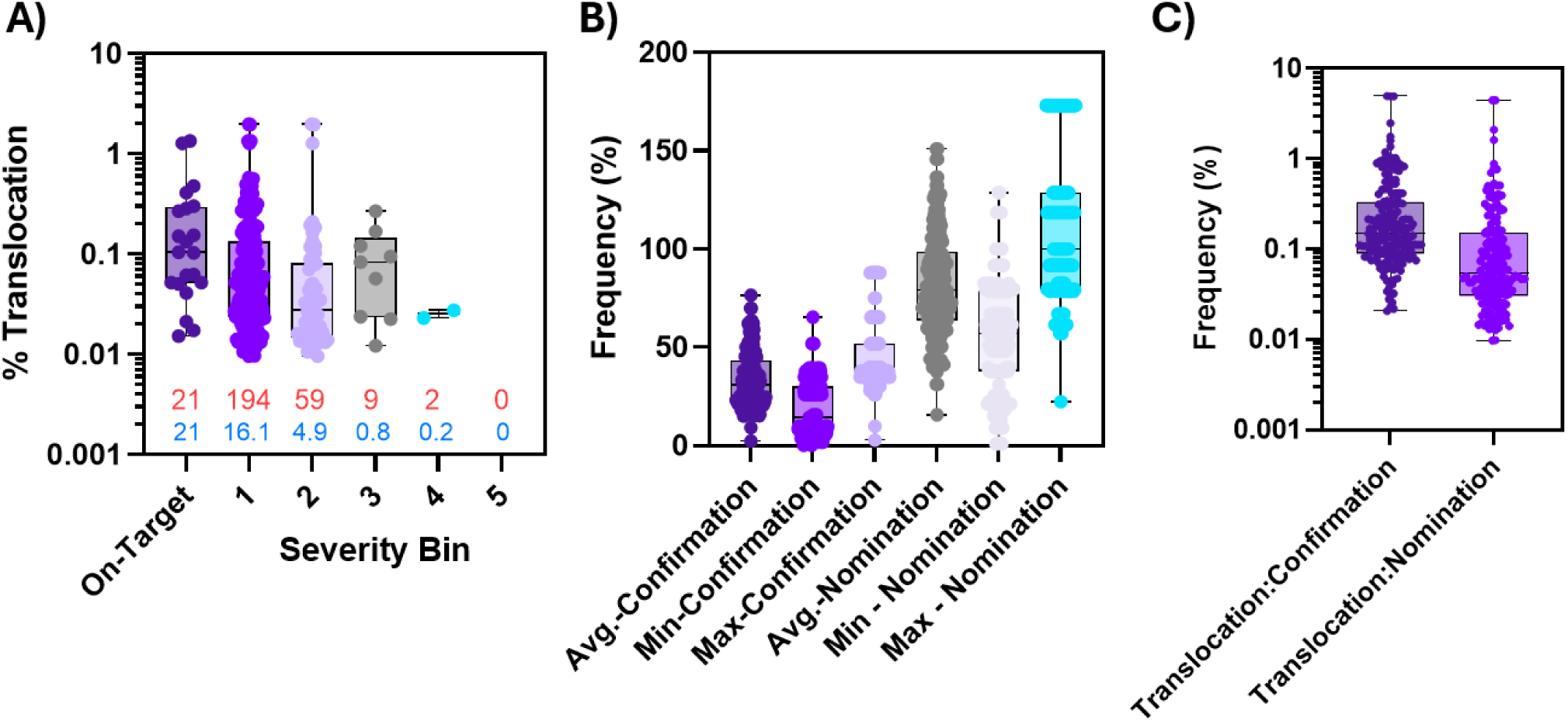
Trends in translocation frequency to other empirical off-target nomination and confirmation features. **A)** Translocation frequency was interrogated relative to the Severity Bin (UNCOVERseq) of different off-targets in addition to the total # of translocation events involving a site (red text) and the average number of translocations per off-target in that Severity Bin group (blue text). **B)** The frequency of nomination (UNCOVERseq; % On-Target UMI Reads) and confirmation (% Indels; rhAmpSeq) was quantified for all significant translocation and the average (Avg.), minimum (Min), and maximum (Max) frequency of each feature calculated between both partners involved in the translocation. **C)** The translocation to confirmation ratio (Translocation:Confirmation) and nomination ratio (Translocation:Nomination) were calculated and plotted

We then evaluated how confirmed indel propensity relates to translocation propensity to gain a better understanding of trends for translocation occurrence. Similar to our observations for nomination propensity, translocation events were generally associated with high frequencies of editing on average between the off-target partners, with an average of 33.2% indel editing between translocated off-target partners (Figure 4B). As was observed with nomination propensity, generally translocations required at least one partner with relatively high quantifiable indel frequencies, with the lowest maximum within a pair being 2.9% indel editing, but an average on 19.65% indel editing (Figure 4B). To attempt to derive a general translation of translocation frequencies to observed average indels, we derived the ratio between the translocation and average indel frequency between partners. On average, the translocation frequency was 0.3% of the average indel frequency between edited partner sites, but it had a range from 0.02% to 4.9% of the indel frequencies between both partners (Figure 4C). This finding supports that translocations are generally a relatively rare proportion of events between two edited partner loci, and that higher indel frequencies between two sites largely leads to higher translocation frequencies.

## Discussion

In this study, we introduce Primer-Anchored Statistical Translocation Analysis (PASTA) as a statistically rigorous and scalable framework for quantifying chromosomal translocations from targeted amplicon sequencing data. By reframing translocation detection as a count-based differential event problem, PASTA leverages mature statistical methodologies originally developed for RNA-seq analysis to achieve performant sensitivity (>75%) and specificity (∼100%) for detecting down to 0.01% translocations for genome editing safety assessment. Furthermore, by testing multiple quantitation algorithms, we show PASTA achieves high quantitative accuracy relative to ddPCR across a range of translocation frequencies representative of real heterogeneous genotoxicity data (0.1% - 10% frequencies). By connecting translocation observations to other empirical datatypes common for genotoxicity assessment, we further elucidate some rough trends that can be used to guide early- and late-safety assessment in identification of translocation likelihood. Importantly, this performance and findings are achieved without requiring specialized library preparation strategies, enabling re-analysis of highly multiplexed amplicon sequencing datasets commonly generated in preclinical genome editing workflows.

Quantitative precision in detecting translocations is an important analytical quality for understanding the genotoxic risk of an event and comparing risk across editing conditions, guide RNAs, or delivery modalities. While previous works in post-hoc amplicon sequencing algorithms have successfully achieved statistical classification of spiked in events down to ∼0.1%, these algorithms had no method for quantitative conversion and relied on absolute count-based reporting of events^24^. Quantitative benchmarking performed in this study to maximize the PASTA algorithm concordance with ddPCR highlights the importance of appropriate normalization strategies when transforming identical NGS count data into biologically interpretable translocation frequencies to avoid systematic errors in this workflow. This mirrors findings in gene editing indel detection algorithms, where identical ground truth frequencies could lead to variable indel frequencies based an algorithm used^16^. Comparatively, we find that the CAST-seq method systematically underestimates translocation frequency, though it is unclear if this is an assay-based or algorithm-based discrepancy. To this end, future work should look more broadly at optimization of quantitative accuracy of different methods through expansion of ground truth datasets that can be easily shared for different workflows directly as gDNA.

DNA DSB induced translocations can occur in variable conformations that may hinder some detection methodologies, with monocentric, dicentric, and acentric events all being possible. In the case of fixed-adapter and random-adapter amplicon sequencing, fixed adapter chemistry is unable to sequence ∼50% of these events, with the specific conformations missed being dependent on the chromosomal location of the initial breaks and on assay orientation. Monocentric translocations are largely more likely to have potential to persist in a population, while acentric and dicentric translocations are often lethal and rapidly lost^37^. While translocations resulting from genome editing have been shown to be possible in all conformations and persist stably, relevant cells such as mESCs and primary T-cells sustaining translocations are often quickly lost over time likely due to fitness impacts^10,11,24,38^. In our work we provide additional support that across individual (on:off_1_) and cumulative (on:off_all_) translocation events, the total translocation burden is often roughly split quantitatively between the different translocation configurations (monocentric, dicentric, acentric) shortly after editing. Thus, both random and fixed adapter chemistry (even with loss of select conformations) still meaningfully provides utility for translocation classification and risk assessment, but with the knowledge that fixed adapter chemistry provides a generally predictable underestimate of the total events and quantitative frequency that are observed for translocation partners deemed significant down to ∼0.01% frequencies. A remaining question here is whether the sensitivity of fixed adapter chemistry is dependent on gDNA isolation time for select events, especially those that generate predominantly acentric/dicentric events that may be more quicky lost in the population. To this end, additional work should be done to understand more comparatively how chemistries may differ on sensitivity with regards to time of genomic DNA isolation.

Analytical sensitivity and specificity are additional translocation detection metrics needed to increase resolution on rare events while minimizing annotation of false positives This relationship is an important consideration for those reviewing diverse analytical approaches, since largely all analytical methods will have some degree of measurable Type I and Type II errors. Through this work, we find that identification of translocations by modeling of biological replication and overdispersed count data through a mature negative binomial framework implemented routinely in count-based RNA-seq analysis produces a performance balance of sensitivity and specificity. Parallel findings to ours can be found in the somatic variant calling field, where similar approaches have been implemented to detect SNPs in variant calling data with similar performance to technology utilizing unique molecular indices (UMIs) in the read structures for error correction^30^. We find here that the limit of detection of 0.01% we annotate largely approaches the actual genomic equivalents in the assay (∼30,000 genomes), leading us to believe that increasing genomic input is one of the most important features for further increasing sensitivity. In contrast, single-sample statistical approaches such as the hypergeometric test exhibited a marked decline in specificity at high coverage, underscoring the importance of replicate-aware and statistically optimized inference for rare event detection. Previously developed amplicon sequencing translocation analysis methods are replicate-unaware and count-based, making them likely prone to these error modes^24^. Furthermore, we find that hypothesis filtering based on minimum counts are necessary default considerations for amplicon-based translocation assessment algorithms since the combination of panel size and background index hopping rapidly expands hypothesis testing into high-dimensional space that can lead to false negatives. These findings align with broader lessons from other areas of genomics, like RNA-seq, where principled statistical modeling and experimental replication are needed to ensure performance in high-dimensional hypothesis testing^25,36^. While we observe high performance metrics in our characterization of PASTA at controlled dilution of a promiscuous gRNA, it is a notable limitation that this analysis only includes translocations present in F/R compatible conformations, and thus will be not completely representative of all occurring events. Future work should focus on expanding the variability of ground truth datasets (panel sizes, translocation frequencies, library preparation chemistries, etc.) with additional orthogonal frequency confirmation to be used as a gold standard set.

While we demonstrate a viable approach for sensitive amplicon sequencing based translocation assessment, alternative technologies also exist for this purpose. Anchored uni-directional preparation methods such as CAST-seq, AMP-seq, and UDiTaS have been able to successfully characterize events down to sub-0.1% frequencies, with UDiTas having additional evidence demonstrating strong linearity in its range of detection^14,23,39^. A distinct advantage of these technologies is that they can additionally identify homology-mediated and off-target-independent translocations from the anchored regions, though these appear to be rare events to date^14,35^. Currently only AMP-seq has demonstrated evidence of being multiplexed to both on- and off-targets to date though, meaning that adoption of many of these workflows for identification of full off-target:off-target translocation interactions is a time-consuming process, though this could likely be implemented through future work. More broadly though, comprehensive metrics on sensitivity, specificity and quantitative precision are still outstanding for many of these assays, and represent areas for additional characterization to better enable translational use in the genotoxicity safety assessment. Comparatively, our PASTA approach largely re-uses existing amplicon sequencing datasets often used for genome editing characterization and scales directly with the multiplexing capabilities of these technologies, while enabling sensitive, specific, and highly linear detection down to 0.01%. Furthermore, we demonstrate that off-target:off-target translocations are relevant events as well, with the likelihood of occurrence largely being directly tied to the nomination and indel frequencies of both sites.

By looking at the biological relationship of translocations to other quantifiable metrics in the genome editing safety assessment process, we were able to support that translocations most commonly require significant indel editing in both translocated partners and largely occur at only a fraction of the events observed by either partner. Previous work from others support this finding, where the identified translocations from gene editing largely involved on-targets and off-targets which had already undergone significant gene editing^10–12,35^. Using these principles may ultimately be helpful for reviewing gene editing safety assessment to guide logic for recommending additional orthogonal methods for quantification or streamline conclusions that a false positive was observed through experimentation. Other work has also investigated and discovered the potential for homology directed repair donors or random chromosomal breaks to facilitate low frequency translocations^14,35^. While some of these events may not be currently captured by this approach, it may be possible by adapting additional primers to sit at commonly identified genomic fragile sites or within the homology directed repair donor itself.

Taken together, these results position PASTA as a practical and relevant solution for chromosomal translocation analysis in translational genome editing applications. By combining statistical rigor, quantitative accuracy, and compatibility with existing amplicon sequencing workflows, PASTA enables detection of rare rearrangements at scales required for modern preclinical safety assessment. Beyond translocation analysis, the framework presented here additionally illustrates the broader potential of adapting well-validated transcriptomic statistical models to other rare-event genomics applications. As genome editing technologies continue to advance toward clinical deployment, such analytically robust and benchmarked methods will be essential for ensuring reproducible, transparent, and trustworthy evaluation of genotoxic risk.

## Methods

### Human cell culture and transfection (K562 and HEK293-Cas9)

K562 (ATCC) and HEK293-Cas9 (ATCC) cells were cultured in Iscove’s Modified Dulbecco’s Medium (IMDM; ATCC) and Eagle’s Minimum Essential Medium (EMEM; ATCC) supplemented with 10% FBS at 37°C with 5% CO_2_. Ribonucleoprotein (RNP) complexes were formed by mixing Alt-R^TM^ *S.p.* Cas9 Nuclease V3 (IDT) or SpyFi^TM^ Cas9 Nuclease (Aldevron) and Alt-R CRISPR-Cas9 sgRNA (gRNA, IDT) and incubating for 20 minutes at room temperature (Molar Ratio: 1:1.2, Cas9:gRNA). For each transfection, 8.0 x 10^5^ cells were washed with 1X phosphate-buffered saline, resuspended in 20 µL of solution SF (Lonza). For K562 cells, RNP complexes at 4 µM were combined with 5 µM of the gRNA (Supplementary Table 1) into the SF solution, while for the HEK293-Cas9 cells, 5 µM gRNA were added to the SF solution. This mixture was transferred into 1 well of a 96-well Nucleocuvette plate (Lonza) and electroporated using program FF-120 (K562) or DS-150 (HEK293-Cas9). Two nucleofections per replicate were performed and each treatment done in triplicate. Following electroporation, cells were transferred to a 6-well plate preheated with either IMDM or EMEM and were incubated at 37°C with 5% CO_2_ for 72 hours. After incubation, genomic DNA (gDNA) was extracted using the Monarch^TM^ Spin gDNA Extraction Kit (New England Biolabs) according to the manufacturer’s instructions, eluted in low-EDTA TE buffer (IDT, 11-05-01-05), and quantified using a NanoDrop 8000 UV-Vis Spectrophotometer (ND-8000-GL).

### Library Preparation

Genomic DNA was extracted from control and genome-edited cells as described above. Libraries for amplicon NGS were prepared using a previously described rhAmpSeq amplification-based method (IDT) using 100 ng of gDNA input^12,40^. Briefly, the first round of PCR was performed using target-specific primers. A second round of PCR was used to incorporate P5 and P7 Illumina adapters to the ends of the amplicons for universal amplification. Libraries were purified using Agencourt AMPure XP system (Beckman Coulter) and quantified by Qubit 1X dsDNA HS Assay kit (Invitrogen) before sequencing on the Illumina NextSeq platform (150-bp paired end reads). Read demultiplexing was performed on the resulting BCL files using Picard v2.18.9 (https://github.com/broadinstitute/picard) IlluminaBasecallsToFastq.

### Computational Analysis – Confirmation

Analysis of the sequencing data to identify confirmed off-target editing at the nominated sites was performed using CRISPAltRations v1.2.1^16^. To classify significant indel editing events, the window for event quantification was centered on the canonical cut site and events quantified utilizing the default window size for Cas9 (8 bp). To determine whether indels found in the sequencing data could result from bona fide off-target cleavage, indels were grouped by location relative to the cut site (prioritizing minimum distance to cut site) followed by fitting counts of events to a negative binominal model with a Wald test for significance in each location bin per off-target using the DESeq2 package within IDT’s OTEasy tool (Schmaljohn et al., Manuscript in Preparation). For classification of indel off-target editing, the tool requires: 1) sufficient read depth for the site (>1,000x) in all replicates, 2) significant edits to occur at or adjacent to the cut site after optimal alignment, 3) the classified cumulative significant edits to exceed 0.01%, 4) the comparison of treatment/control samples at the site to have a significant adjusted p-value (p < 0.05), and 5) an average coverage frequency of at least 5x the ascribed cumulative frequency observed (e.g., for 0.1% editing, at least 5,000x coverage).

### Computational Analysis – Translocations

To quantify translocations from editing, Primer Anchored Statistical Translocation Analysis (PASTA) was used. This tool was created as dockerized workflow with a high-level workflow as described (Supplementary Figure 1). Briefly, the tool expects the following as inputs: 1) a configuration file in YAML format specifying the locations of all files and parameters to be used, 2) unaligned BAM format sequencing files (uBAM) consisting of paired end sequencing data (e.g. R1/R2 2×150 sequencing), 3) a BED format file specifying the coordinates of all amplicons in the multiplexed panel, and 4) an output directory to put results. The tool begins by using the assay coordinate BED file on a configurable reference genome of interest and extracting the 34 nucleotides 5’ and 3’ of the assay coordinates using pyfaidx^41^ (v0.9.0.3) as assumed primer regions. Following this, primers are identified in R1/R2 reads within the uBAM file input using fg-idprimer (https://github.com/fulcrumgenomics/fg-idprimer; -k=6, -K=8, -S=5, --max-mismatch-rate=0.07) to generate a count matrix of each read’s R1/R2 primer associations. To enable variable downsampling of events in the post-primer identification file, we implemented an optional downsampling methodology off of the count matrix that enables downsampling each assay randomly down to set values in a seeded manner. Following this, count matrices are merged in a analysis-specific manner using pandas^42^ (v2.3.3). Prior to statistical testing, we exposed a flag to filter unique events based on average treatment frequency count to remove invalid hypotheses for over-expansion of post-hoc correction test evaluation (default = 2 reads). Then, treatment/control pairs had their counts paired and primer count frequencies subjected to either: A) one-tailed hypergeometric test with Benjamini-Hochberg correction (statsmodel v0.15.0; default settings), also referred to as the ‘hypergeometric’ method or, B) a negative binomial model followed by Wald-test and Benjamini-Hochberg correction as implemented in the python DESeq2 package^43^ (pydeseq2; v0.5.3) to calculate an adjusted p-value (p-adj). Translocations were significant if: 1) the estimated translocation frequency exceeds 0.005%, 2) the translocation has a significant adjusted p-value (p adj < 0.01), 3) the background frequency of the event is quantified to be less than 1% in unedited samples. Following this, events that were classified as a translocation had the translocation frequency (P) calculated using the following optimized default equation (“mean”):

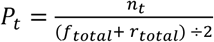

Where ‘n’ is equal to the count of the unexpected primer pair of interest, ‘t’ is the significant translocation being interpreted, ‘f’ is the total count of the shared forward primer events excluding the count participating in the ‘n’ translocation event, and ‘r’ is the total count of shared reverse primer events excluding the count participating in the ‘n’ translocation event. The translocation frequency is then adjusted by the background level frequency in the control by subtracting any translocation frequency observed in the control sample from the treatment frequency. Additional calculation equations for translocation frequency are optionally available and detailed in Supplementary Figure 2.

## Data and Code Availability

The code for executing PASTA is available for download upon request **exclusively for non-commercial research purposes** through DockerHub by messaging the authors (; https://hub.docker.com/repository/docker/idtdna/pasta). Commercial use of the application requires explicit consent from the authors which can be requested by messaging. Re-analyzed datasets from previous publications can be found in their respective publications and NCBI SRA database project PRJNA1380439 and PRJNA1337486. Sequencing data generated in this study have been deposited in the SRA database under accession code PRJNA1496969.

## Disclaimers and Conflicts of Interest

Products and tools supplied in this manuscript by IDT are for research use only and not intended for diagnostic or therapeutic purposes. Purchaser and/or user are solely responsible for all decisions regarding the use of these products and any associated regulatory or legal obligations. For informational use only. The data provided are for informational use only and should not be used as the sole basis for any critical decision making. The data generated are based on assay procedures that have not undergone full validation: formal design and development activities are on-going. M.S., G.K., C.B.., O.U., E.S., T.O., K.J.K.., A.S.P., G.R., and A.M.J. are employees of IDT, which offers reagents and services for sale similar to some described in the manuscript

---

**Supplementary Figure 1.**
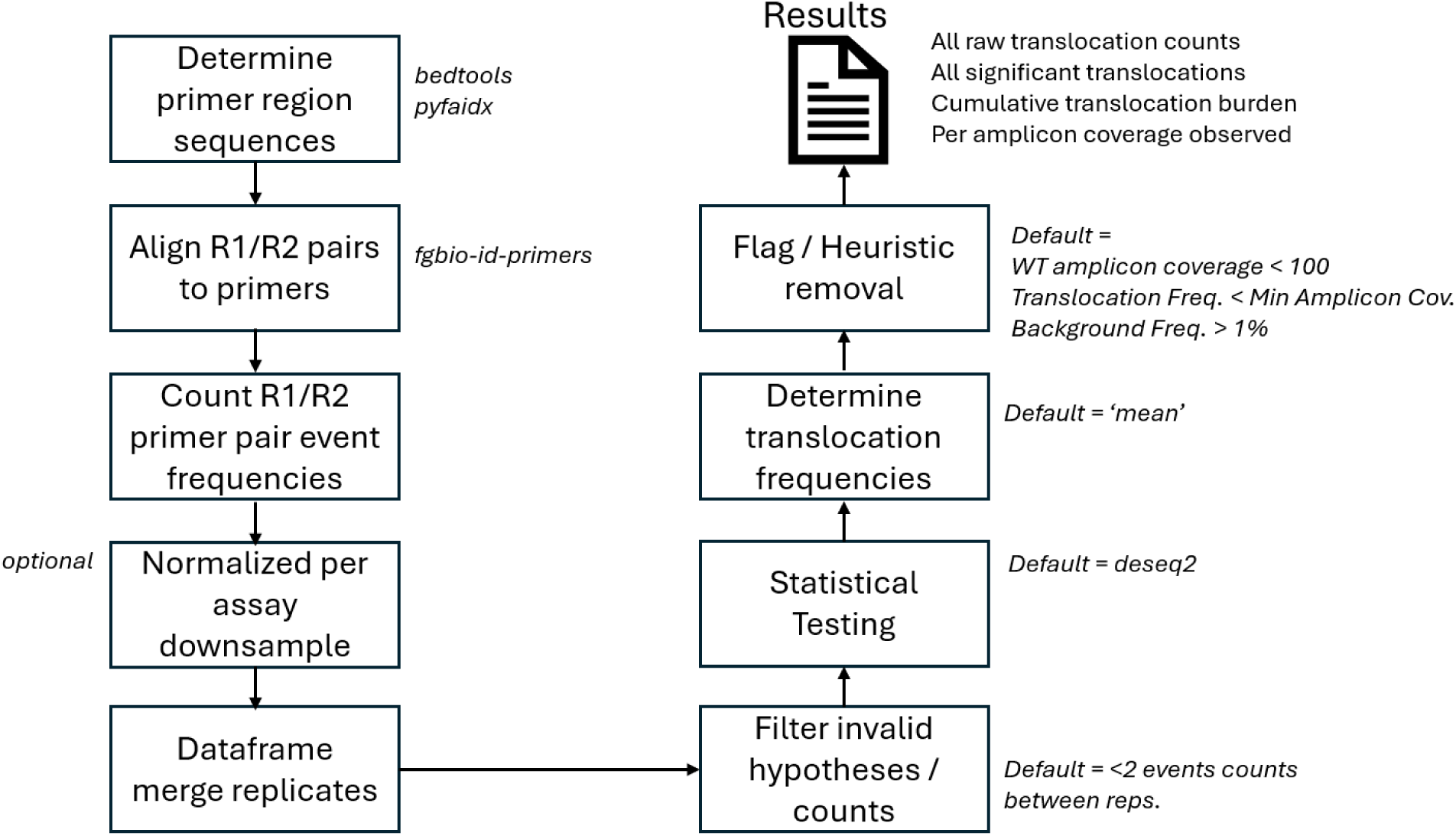
Diagram on PASTA algorithm and defaults. Depicted is the algorithm flow from which 1) raw sequencing files (FASTQ) and 2) amplicon BED files are used to determine the primer region sequences, align the reads to, and then perform event counting, filters, and statistical testing to determine significant translocations that pass criteria for variable heuristic flags.

**Supplementary Figure 2.**
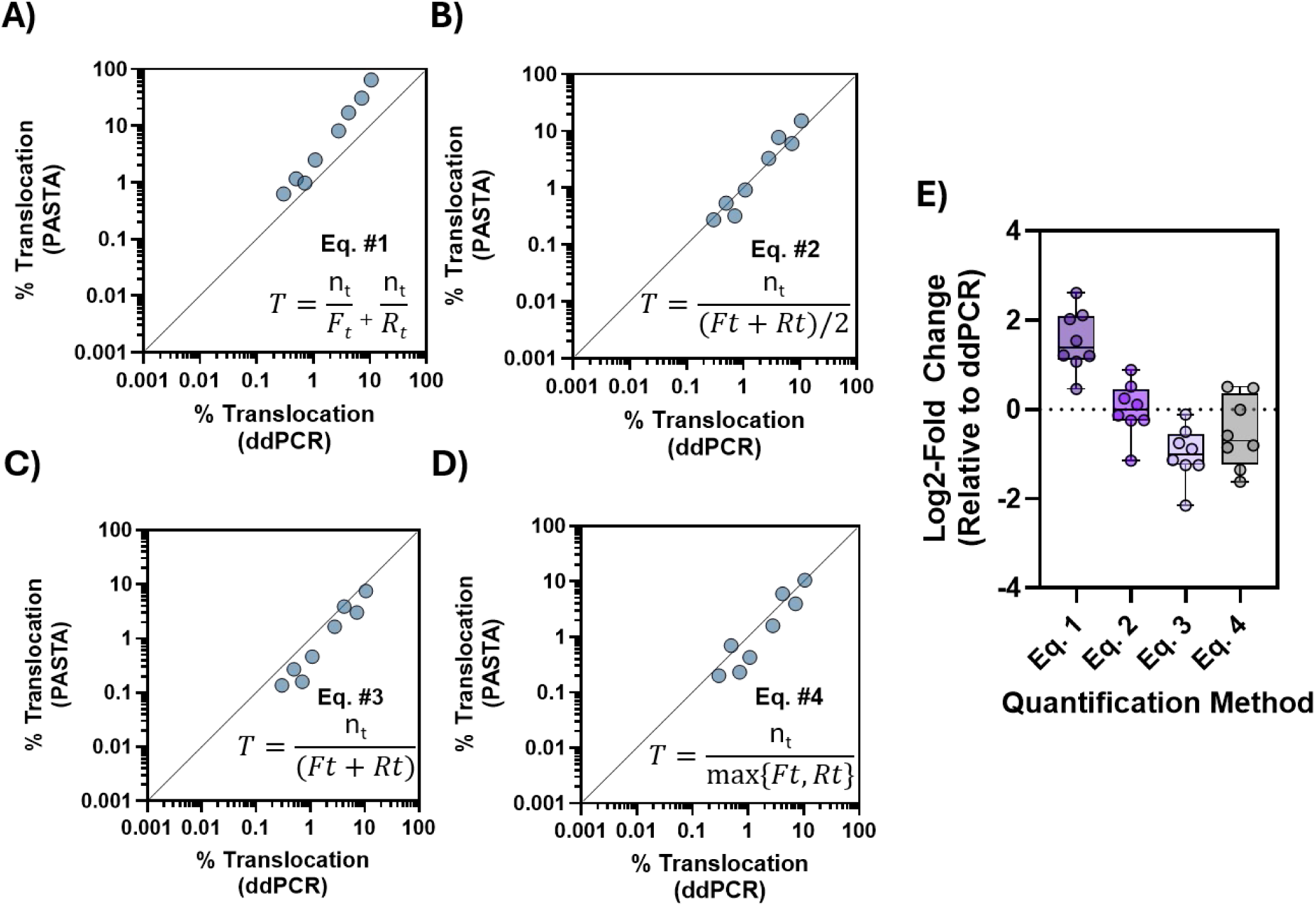
Effect of PASTA Translocation Calculation Method on Quantitative Precision. Four different calculations were tested for deriving translocation frequency values from amplicon sequencing data. Equation is shown in the bottom right for **A)** Equation #1 **B)** Equation #2 **C)** Equation #3 **D)** Equation #4. Variables are as follows: T; Translocation frequency, n_t_; number of translocation events, F_t_; total forward primer events, R_t_; total reverse primer events. **E)** For all equations the calculated fold change from ddPCR was calculated and shown as a box plot.

**Supplementary Figure 3.**
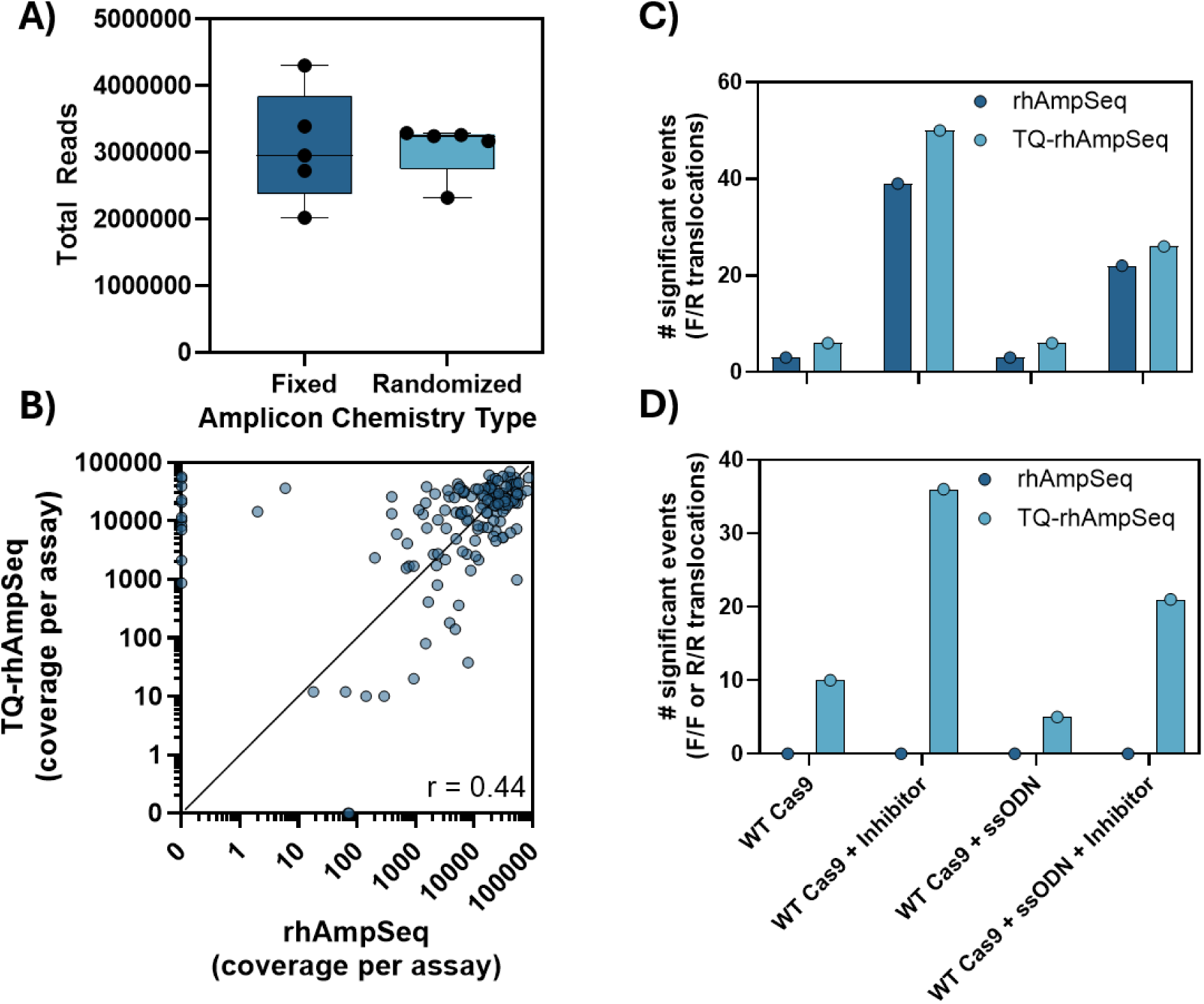
PASTA detection of fixed and randomized adapter amplicon events. **A)** The total number of reads for fixed adapter amplicon chemistry (rhAmpSeq) and randomized adapter chemistry (TQ-rhAmpSeq) was quantified along with **B)** per assay coverage and Spearman r between the two datasets. **C)** The number of **s**ignificant F/R primer translocations and **D)** F/F and R/R primer translocations were quantified in gDNA from samples edited with a CCR5 gRNA with both a fixed adapter chemistry (rhAmpSeq) and a randomized adapter chemistry (TQ-rhAmpSeq). F/F and R/R translocations should only be detectable in randomized adapter chemistry.

**Supplementary Figure 4.**
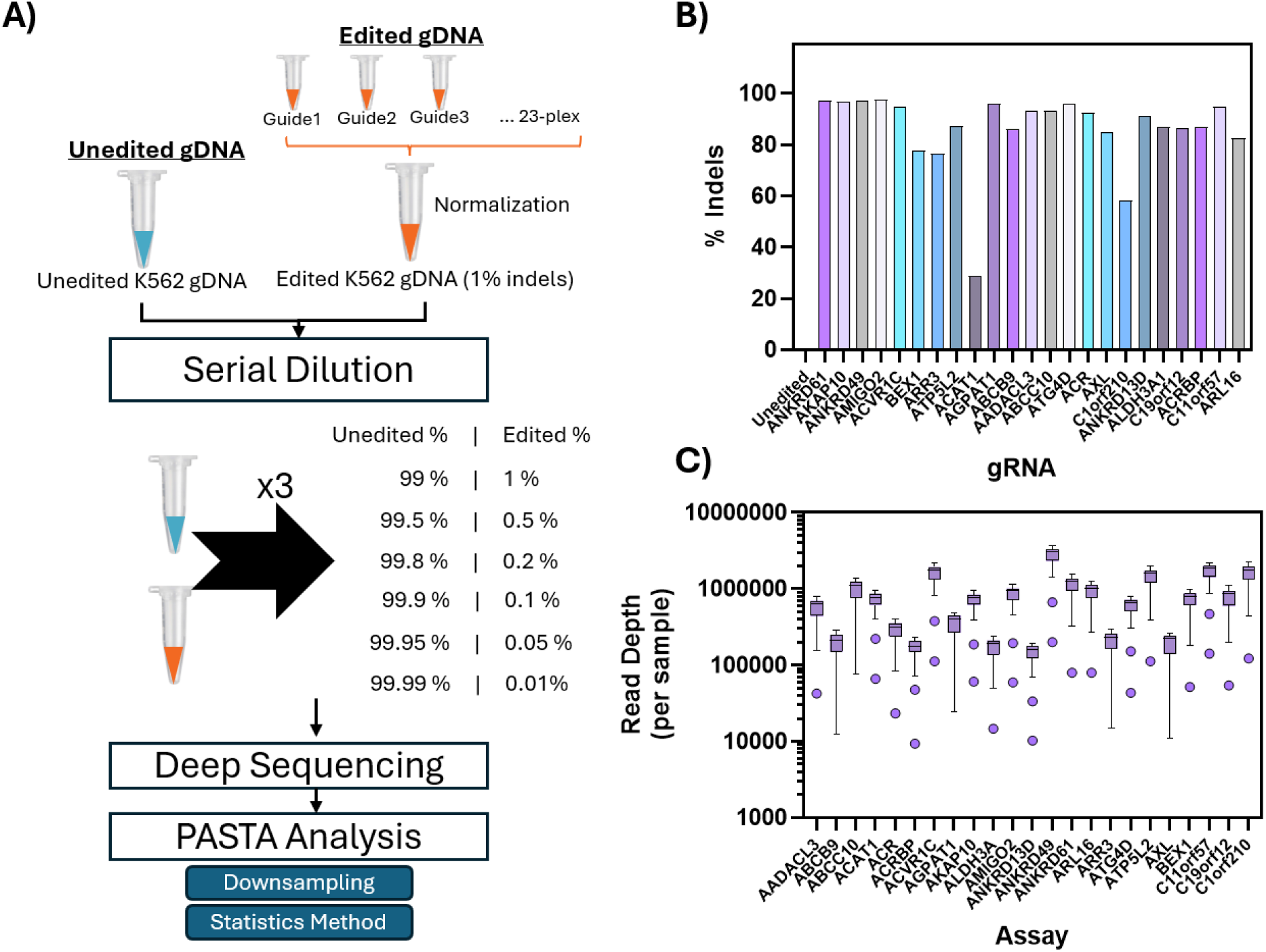
Initial characterization of specificity dataset. **A)** General workflow shown for creating the specificity dataset and testing the effect of read depth and statistics methods on results. **B)** The indel frequencies of the undiluted samples were characterized compared to an unedited control in K562 cells. **C)** Following sequencing of samples with a twenty-three multiplexed primer set of genomic targets that were unrelated to one another (n=24 samples; 3 replicates per dilution) the read depth per site per sample was quantified.

**Supplementary Figure 5.**
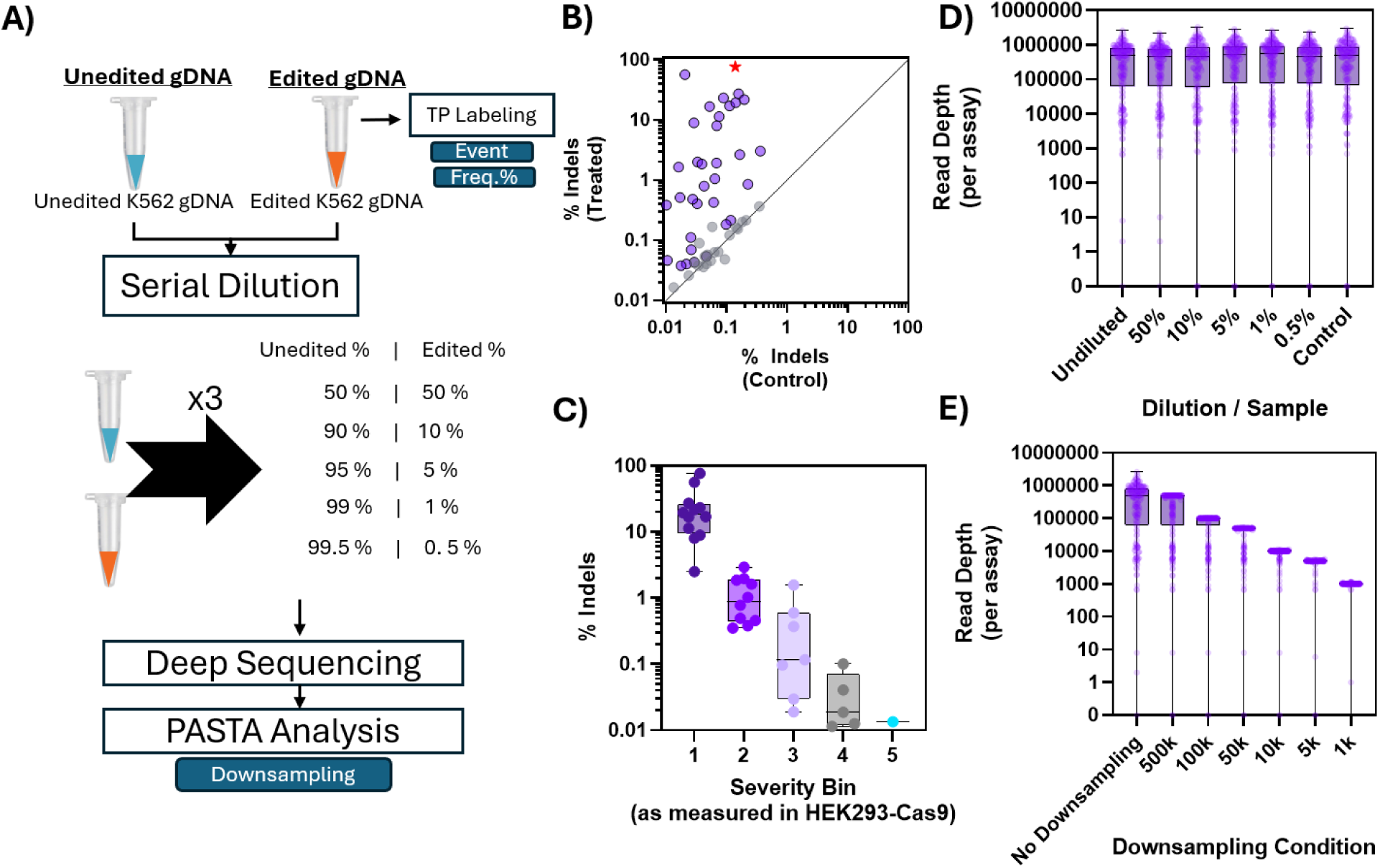
Characterization of the sensitivity translocation dataset. **A)** Workflow description for generating the sensitivity dataset. Briefly, cells were edited with the LAG3 site 9 gRNA and had high confidence translocations characterized in terms of their identity and frequency before being subjected to a serial dilution and analysis using PASTA. **B)** Significantly edited sites were identified and **C)** had the off-target Severity Bin they belong to annotated (as in Kinney et al. 2026). **D)** Read depth of the different assays was quantified per dilution level both pre and **E)** post-downsampling of reads.

**Supplementary Figure 6.**
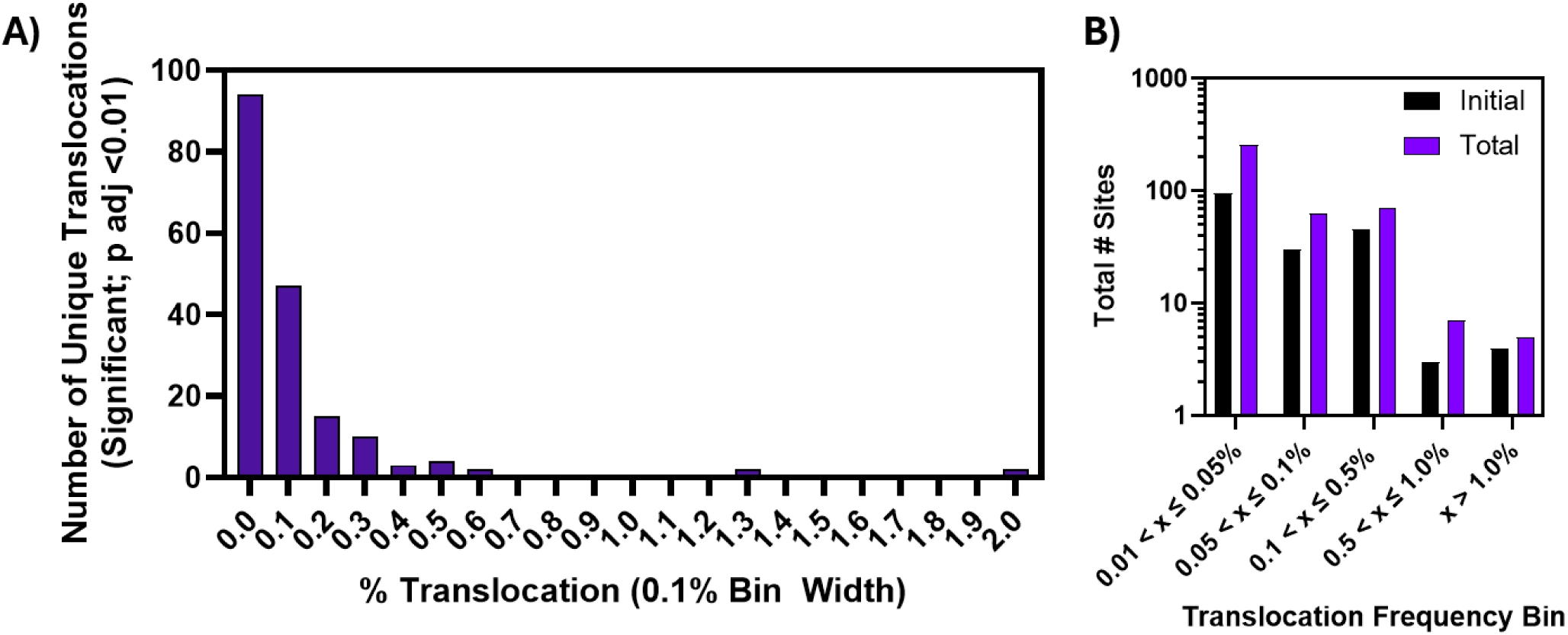
Characterization of translocations frequencies in sensitivity dataset. **A)** Quantification of the average translocation frequency of significant translocations in the undiluted LAG3 site 9 edited samples (n = 3 treatment / control pairs). Frequency was binned in 0.1% translocation intervals and is presented as a histogram. **B)** The number of different true positive sites in the undiluted sample was binned in 0.5% frequency bins and the total # of sites in each bin was quantified both before and after accounting for frequencies being represented or repeated throughout the dilution series.

**Supplementary Figure 7.**
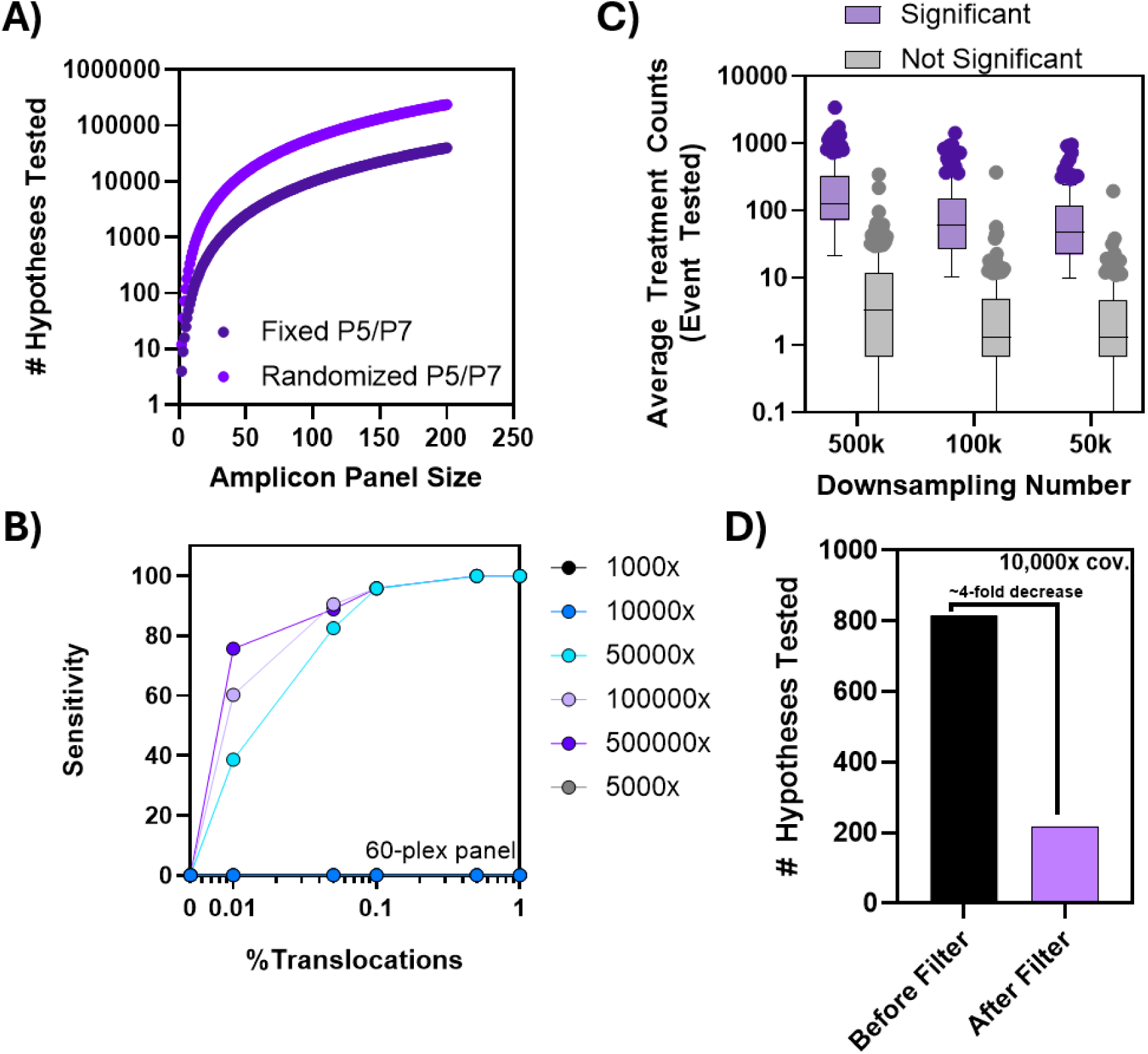
Multiple hypothesis testing issues in significance testing for translocations. **A)** In silico quantification of the number of hypotheses being tested as panel sizes increase for both fixed and random adapter chemistry up to a 200plex. **B)** Sensitivity performance of the LAG3 site 9 60-plex panel at variable read depth confirms performance issues as read depth decreases. **C)** The average count of events in treatment samples measured of both significant and not significant events in the 60-plex panel shows clear count separation between groups. **D)** To measure hypothesis testing numbers with real data, the number of hypotheses tested was measured before and after implementation of minimum average treatment count filter (requirement of n > 2 counts).

